# Novel Loss-of-Function Mutations in *COCH* Cause Autosomal Recessive Nonsyndromic Deafness

**DOI:** 10.1101/2020.04.30.071134

**Authors:** Kevin T Booth, Amama Ghaffar, Muhammad Rashid, Luke T Hovey, Mureed Hussain, Kathy Frees, Erika M Renkes, Carla J Nishimura, Mohsin Shahzad, Richard J Smith, Zubair Ahmed, Hela Azaiez, Saima Riazuddin

## Abstract

*COCH* is the most abundantly expressed gene in the cochlea. Unsurprisingly, mutations in *COCH* underly deafness in mice and humans. Two forms of deafness are linked to mutations in *COCH*, the well-established autosomal dominant nonsyndromic hearing loss, with or without vestibular dysfunction (DFNA9) via a gain-of-function/dominant-negative mechanism, and more recently autosomal recessive nonsyndromic hearing loss (DFNB110) via nonsense variants. Using a combination of targeted gene panels, exome sequencing and functional studies, we identified four novel pathogenic variants (two nonsense variants, one missense and one inframe deletion) in *COCH* as the cause of autosomal recessive hearing loss in a multi-ethnic cohort. To investigate whether the non-truncating variants exert their effect via a loss-of-function mechanism, we used mini-gene splicing assays. Our data showed both the missense and inframe deletion variants altered RNA-splicing by creating an exon splicing silencer and abolishing an exon splicing enhancer, respectively. Both variants create frameshifts and are predicted to result in a null allele. This study confirms the involvement of loss-of-function mutations in *COCH* in autosomal recessive nonsyndromic hearing loss, expands the mutational landscape of DFNB110 to include coding variants that alter RNA-splicing, and highlights the need to investigate the effect of coding variants on RNA-splicing.

## Introduction

The harmonious interplay between many proteins is required for normal auditory function. Mutations in genes encoding these proteins give rise to hearing loss. Each of these genes has a distinct genomic and mutational signatures^1^. For some genes, variant location and variant type correlate with specific phenotypic outcomes as a consequence of the pathogenic mechanism at play; such is the case with the *COCH* gene^2–4^. Cochlin, the protein product of the *COCH* gene, is a large secreted extracellular protein that contains an N-terminus LCCL domain and two von Willebrand factor A-like domains (vWFA1 and vWFA2). It is the most abundantly detected protein in the inner ear^5,6^ and plays a well-established role in post-lingual progressive autosomal dominant nonsyndromic hearing loss (ADNSHL), sometimes accompanied by vestibular dysfunction, at the DFNA9 locus^4,6–8^ (Figure 1E). DFNA9 deafness occurs via multiple mechanisms depending on mutation location^4,8–10^. Mutations in the LCCL domain typically cause misfolding and defective multimerization of cochlin^4^. Patients with these mutations typically have later onset hearing loss that is accompanied by vestibular dysfunction^4^. In contrast, mutations in the vWFA domains typically result in secretion failure and aggregate formation^4^. Patients with these mutations report earlier onset hearing loss without vestibular dysfunction^4^. While the toxic gain of function/dominant-negative effects of mutant cochlin on the auditory system and the underlying pathology that results in hearing loss have been well studied^4^, little is known about the consequence and underlying pathology from loss of cochlin.

**Figure 1.**
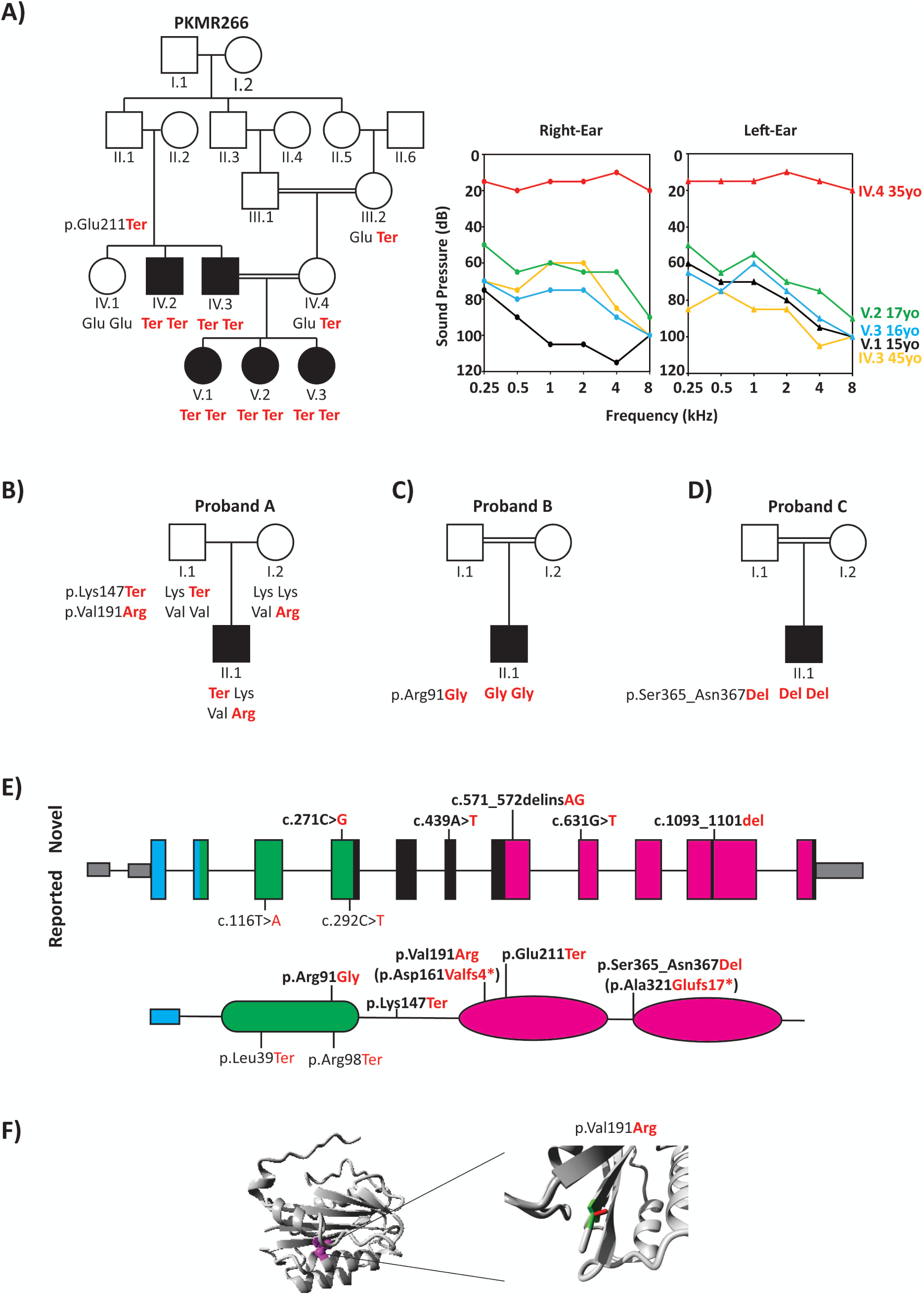
Pedigrees, audiograms, and *COCH* gene and protein schematic. **(A)** Segregation of p.Glu211Ter and audiograms for family PKMR266. Audiograms were obtained with air conduction with the frequency range of 250–8000Hz. **(B-D)** Genotypes of Probands A, B, and C and their available family members. **(E)** Human *COCH* (NM_004086.2) has 12 exons, of which the first exon is not coding (grey). Variants identified in this study denoted in bold. The COCH protein encodes a 550 amino acid peptide consisting of a signal peptide (blue), an LCCL domain (green), and two VWFA domains (pink). Novel variants identified in this study (top) and previously reported DFNB110 variants (bottom). Square brackets represent the predicted effect of the splicing variants. **(F)** Three-dimensional modeling showing an overview of protein in ribbon presentation (p.Val191 highlighted in magenta) and locally zoomed overlay of wildtype p.Val191 (green) and mutant p.Arg191 (red).

Recently, two families with autosomal recessive nonsyndromic hearing loss (ARNSHL) at the DFNB110 locus have been described segregating nonsense variants in *COCH*^2,3^, suggesting expression of COCH is required for proper auditory function. Here we expand on these findings to report four novel mutations associated with DFNB110 deafness and demonstrate the involvement of missense and inframe variants in mRNA splicing. Our findings further support loss-of-function as a third pathogenic mechanism associated with *COCH*-related hearing loss in humans.

## Methods

### Subjects

One Pakistani (PKMR266) family, and three probands of European (Proband A), Middle-Eastern (Proband B), and unknown ethnicity (Proband C) segregating ARNSHL were ascertained for this study. Affected individuals underwent clinical examination and pure tone audiometry to measure hearing thresholds at 0.25, 0.5, 1, 2, 3, 4 and 8 kHz. After obtaining written informed consent to participate in this study, blood samples were obtained from all affected and unaffected family members and genomic DNA was extracted. The human research Institutional Review Boards approved all procedures at the University of Iowa, Iowa City, Iowa, USA, University of Maryland, School of Medicine, Baltimore, USA, and Shaheed Zulfiqar Ali Bhutto Medical University, Islamabad, Pakistan

### Next-Generation Sequencing and Bioinformatic Analysis

Probands A-C underwent comprehensive genetic testing to screen all known genes implicated in NSHL, common NSHL mimics, and common syndromic forms of deafness using the OtoSCOPE^®^ panel as described^11–14^. Similarly, a proband from family PKMR266 underwent genetic testing as described^15,16^. Bioinformatic and variant analysis were also completed as described^13–15^. In brief, after mapping of raw sequence reads and variant calling, variants were annotated and filtered based on quality (depth >5 and call quality >30), minor allele frequency (MAF) <2% in Genome Aggregation Database (gnomAD)^17^ and variant effect (missense, nonsense, indel or splice-site). Retained variants were prioritized based on their conservation (GERP and PhyloP) and predicted deleteriousness (SIFT, PolyPhen2, MutationTaster, LRT, and the Combined Annotation Dependent Depletion (CADD))^18,19^. Variant effect on splicing was assessed using Human Splicing Finder (HSF)^20^. Samples also underwent copy number variant (CNV) analysis. Segregation analysis was carried out on available family members using Sanger sequencing.

### *In vitro* splicing analysis

*In vitro* splicing assays were performed as described^13,14,21^. *COCH* (NM_004086) gene-specific primers were used to amplify exon 5, 7, 8 and 11 using patient genomic DNA and control DNA to obtain the mutant and wildtype allele, respectively. The amplicons were then ligated into the pET01 Exontrap vector (MoBiTec, Goettingen, Germany). Colonies were selected and grown, and plasmid DNA was harvested using the ZymoPure Plasmid Midiprep Kit (ZYMO Research, Irvine, California, USA). After sequence confirmation, wildtype and mutant minigenes were transfected in triplicate into COS7, HEK293, and MDCK cells, and total RNA was extracted 36 hours post-transfection using the Quick-RNA MiniPrep Plus kit (ZYMO Research). Using a primer specific to the 3′ native exon of the pET01 vector, cDNA was synthesized using AMV Reverse Transcriptase (New England BioLabs). After PCR amplification, products were visualized on a 1.5% agarose gel, extracted and then sequenced.

### Molecular modeling and Thermodynamic Predictions

Molecular modeling of COCH is based on the protein structure PDB: 4IGI. The model was built using Yasara and WHAT IF Twinset^22^ using the project HOPE server (http://www.cmbi.ru.nl/hope/). Thermodynamic predictions were calculated using STRUM^23^ server (https://zhanglab.ccmb.med.umich.edu/STRUM/).

## Results

### Subjects and Variant Identification

We ascertained a consanguineous Pakistani family and three probands of European (Proband A), Middle-Eastern (Proband B), and unknown ethnicity (Proband C) with ARSNHL (Figure 1A-D). Hearing loss in family PKMR266 is prelingual moderate-to-profound (Figure 1A). Audiograms could not be obtained for probands A-C, however clinical description and history were provided. Proband A is an 11-year-old male with congenital and moderate hearing loss. Proband B is a 3-year-old male with congenital down-sloping mild-to-severe hearing loss. Finally, Proband C is a 15-year-old male with prelingual down-sloping mild hearing loss. Families for probands B and C also reported consanguinity.

Combination of OtoSCOPE^®^ panel and NGS was used to identify underlying genetic cause of hearing loss segregating in these affected individuals. Genetic variant filtering for MAF, quality, effect, and recessive or X-linked inheritance yielded candidate variants in the *COCH* gene for all probands. In Family PKMR266, a homozygous nonsense variant [c.631G>T; p.(Glu211Ter)] in exon 9 was identified, which segregated with the hearing loss phenotype in 5 affected individuals spanning two generations. In Proband A, we identified two variants in *COCH*: a nonsense variant (c.439A>T; p.Lys147Ter) and a dinucleotide change (c.571_572delinsAG; p.Val191Arg), in exons 7 and 8, respectively (Figure 1E). Proband B carries a homozygous missense variant (c.271C>G; p.Arg91Gly) in exon 5, and in Proband C a homozygous inframe deletion (c.1093_1101del; p.Ser365_Asn367del) in exon 11 was identified. All identified variants are novel (absent from gnomAD) or ultra-rare (Table 1) and occur in conserved residues (Table 1). Variants p.Arg91Gly and p.Val191Arg are predicted to be deleterious by computational tools (Table 1). No CNVs were detected in patients screened by NGS. We also identified two variants in *LOXHD1* in Proband B (Supp Table 1). Both variants are rare and have conflicting scores as to conservation and predicted deleteriousness.

**Table 1.** Novel and Previously reported DFNB110 variants.

| ID | Ethnicity | g.DNA | c.DNA | Protein | Max<br>gnomAD<br>(%) | GERP | phyloP | SIFT | PP2 | MT | LRT | CADD | $\Delta\Delta G$ | Ref |
| --- | --- | --- | --- | --- | --- | --- | --- | --- | --- | --- | --- | --- | --- | --- |
| 9600236 | Iranian | 14:g.31346811T>A | c.116T>A | p.Leu39Ter | 0 | 5.67 | 1.03 | - | - | A | N | 36 | - | <sup>3</sup> |
| Proband B | Middle Eastern | 14:g.31348048C>G | c.271C>G | p.Arg91Gly | 0 | 2.47 | 0.932 | T | D | D | D | 19.64 | -2.77 | * |
| AI-family | Belgian/<br>Moroccan | 14:g.31348069C>T | c.292C>T | p.Arg98Ter | 0.008 | 3.01 | 0.932 | - | - | A | N | 37 | - | <sup>2</sup> |
| Proband A | European | 14:g.31349660A>T | c.439A>T | p.Lys147Ter | 0 | 5.82 | 1.2 | - | - | A | D | 41 | - | * |
|  |  | 14:g.31349882_31349883GT>AG | c.571_572delinsAG | p.Val191Arg | 0 | - | - | D | - | - | - | - | -0.02 | * |
| PKMR266 | Pakistani | 14:g.31353760G>T | c.631G>T | p.Glu211Ter | 0.003 | 5.92 | 1.05 | - | - | A | D | 43 | - | * |
| Proband C | Unknown | 14:g.31355131_31355139del | c.1093_1101del | p.Ser365_Asn367del | 0 | - | - | - | - | - | - | - | - | * |

Nucleotide numbering: the A of the ATG translation initiation site is noted as +1 using transcript NM\_004086.2 of *COCH*. PP2, PolyPhen2; MT, MutationTaster; D, predicted Damaging or Deleterious; -, data not available; T, Tolerated; A, disease automatic; An “\*” indicates variant identified in this study.

### Computational and in vitro splicing analysis

Since the previously described DFNB110-causing mutations were all truncating loss of function variants, we sought to explore the effects of the missense and inframe variants found in Probands A-C on RNA splicing. Variants c.271C>G, c.571_572delinsAG, and c.1093_1101del, were computationally predicted to affect splicing motifs and impact splicing by various mechanisms. The c.271C>G variant is predicted by HSF to alter an exonic splicing enhancer (ESE), create an exonic splicing silencer (ESS), and create a cryptic donor site. RT-PCR of cells transfected with either wildtype or mutant minigenes revealed no alterations in splicing between the wildtype and mutant (Figure Sup 1). The p.Arg91 residue is located in the LCCL domain (Figure 1E). Computational 3D modeling was not possible due to a lack of structure resolution. We could, however, calculate the change in folding energy between the wildtype and mutant^23^. The calculated difference, −2.77, strongly suggests the p.Arg91Gly has an impact on protein stability.

The loss of nucleotides c.1093-1101 in exon 11 is predicted to break the binding motifs for the serine/arginine-rich splicing factors 1 and 2 (SRSF1 and SRSF2). Visualization and sequencing of splicing products from wildtype and mutant minigenes revealed the inclusion of exon 11 in the wildtype but not in the mutant (Figure 2A). Loss of exon 11 creates a shift in the reading frame and result in a truncated protein product (p.Ala321Glufs17*).

**Figure 2.**
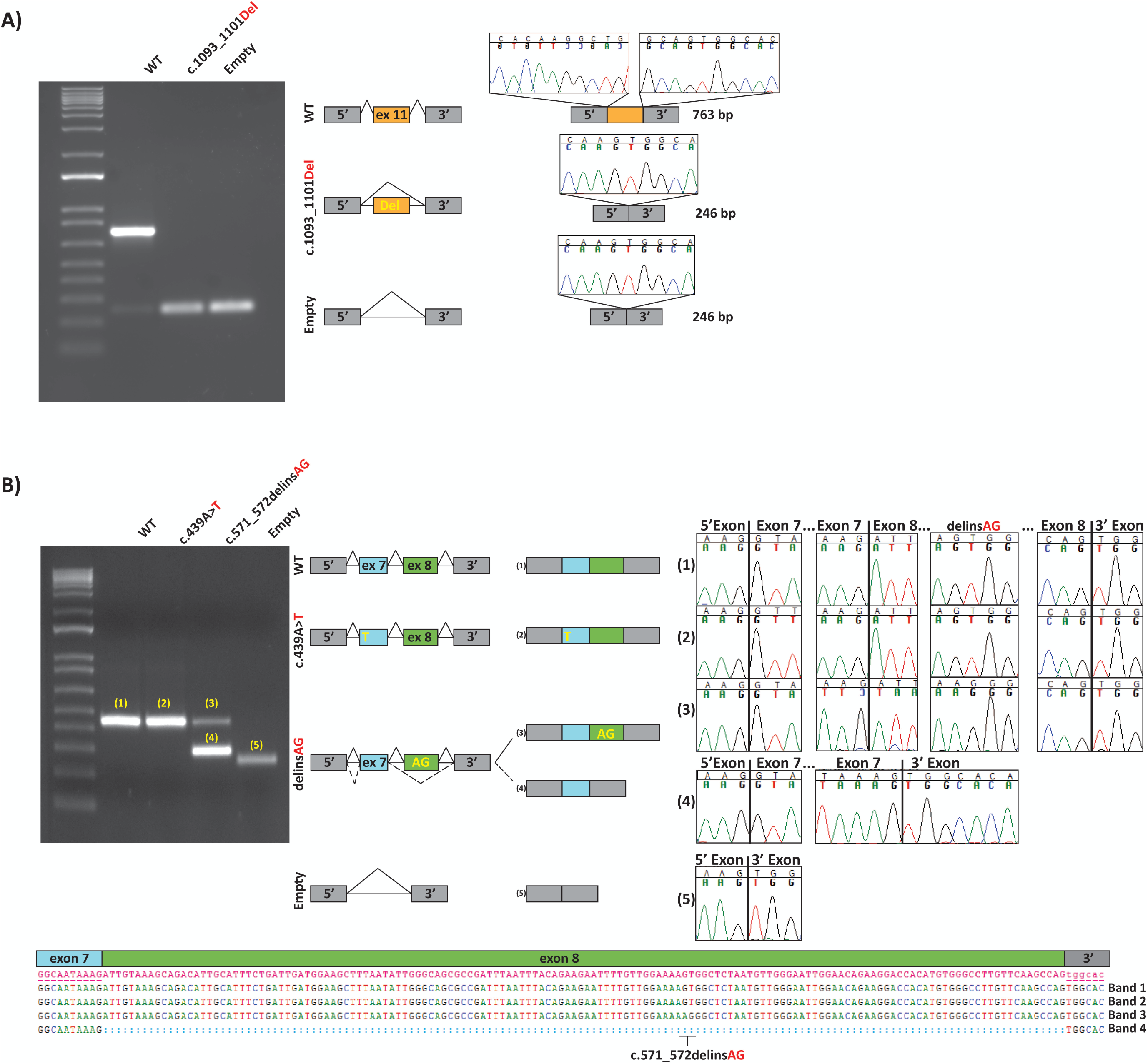
Minigene splicing assays. Gel electrophoresis and sequence chromatograms of wildtype, empty pET01 vector, and the c.1093_1101del (A) and c.571_572delinsAG (B). **(A)** The c.1093_1101del variant abolishes an exonic splicing enhancer in exon 11, resulting in complete exon skipping confirmed by Sanger sequencing. **(B)** The dinucleotide c.571_572delinsAG variant produced two bands. The top band (#3) corresponds to a normally spliced product and migrates in parallel with the wildtype and the c.439A>T variant. The smaller band (#4) corresponds to the aberrant splicing of exon 8. Sequence chromatograms show the read through at each exon junction, and sequence alignment shows the deletion of exon 8 in-band #4 and the dinucleotide change in band #3.

Finally, the c.571_572delinsAG alteration is predicted to activate 2 cryptic acceptor sites and create an ESS. Visualization of the spliced products for the mutant minigene carrying the c.571_572delinsAG revealed two bands at ∼440bp and ∼300bp (Figure 2B). Sequencing showed the band corresponding to ∼440 bp contained both exon 7 and 8, whereas the smaller band (∼300 bp) revealed skipping of exon 8 (Figure 2B). The loss of exon 8 in the mRNA shifts the reading frame resulting in a premature stop codon (p.Asp161Valfs4*). Since proband A DNA was used to create the minigene we used the *in trans* nonsense c.439A>T variant in exon 7 as control for splicing efficiency. Minigenes carrying the c.439A>T variant revealed a single band at ∼440 bp, identical to the wildtype; subsequent sequencing revealed the presence of the c.439A>T variant but no alterations to wildtype splicing (Figure 2B).

## Discussion

We used a combination of targeted genomic enrichment and massively parallel sequencing to implicate *COCH* as the causal gene in one large and three small families segregating ARNSHL. Among the identified *COCH* pathogenic variants, two are nonsense, one a dinucleotide change, one a missense variant, and one is an inframe deletion. Of these variants, three are novel (not reported in the Genome Aggregation Database, gnomAD; Table 1). To date, only two nonsense variants have been linked to *COCH*-related ARNSHL (Figure 1E, Table 1).

In humans, *COCH* is the most abundantly expressed gene in the cochlea^5,24^. The twelve-exon gene encodes a 550 amino acid Cochlin protein consisting of a signal peptide, an LCCL domain, and two vWFA domains (Figure 1E). Unlike most genes in the inner ear linked to deafness, *COCH* is highly expressed in the fibrocytes of the cochlea and not in the mechanosensitive hair cells^25–27^. After translation, Cochlin is secreted from the fibrocytes. While its exact function is unknown, it has been suggested that Cochlin might play an essential structural role in organizing and stabilizing the extracellular matrix via its vWFA domains. More recently, it has been shown that the LCCL domain is critical for the immune response in the inner ear^28^. Its role in hearing loss implicates Cochlin as a protein required for proper auditory function in humans.

The ultra-rare nonsense variant c.631G>T (p.(Glu211Ter)) segregating with the moderate-to-severe deafness in family PKMR266 (Figure 1A) occurs in exon 9 (Figure 1E). Given that the created termination codon is over 50 base pairs away from the last exon-exon junction, we predict that nonsense-mediated decay (NMD) will occur^29,30^, leading to a null allele. If the allele carrying this variant was to escape NMD, the translated truncated protein product would be missing ∼50% of the protein (Figure 1E), rendering the protein nonfunctional. The unaffected mother (IV:4) is heterozygous for the p.(Glu211Ter) variant and has normal hearing at the age of 35. These findings show that haploinsufficiency is not a pathogenic mechanism for *COCH*-related hearing loss^2,3,31^.

Novel compound heterozygous variants ((c.439A>T; p.Lys147Ter) and (c.571_572delinsAG; p.Val191Arg)) were identified in Proband A. Similar to the p.Glu211Ter variant identified in family PKMR266, p.Lys147Ter variant is predicted to produce a null allele via the NMD pathway^29,30^. The dinucleotide change c.571_572delinsAG replaces a GT with an AG and results in a single amino acid substitution valine-to-arginine at the highly conserved residue 191 (Figure 1E and Supp Figure 2). Molecular modeling for this variant revealed that residue 191 is buried in the core of the vWFA1 domain (Figure 1F). The change from the neutral valine to the positively charged and bulkier arginine could disable the protein and alter folding as shown by the negative change in free energy (Table 1) predicting the mutant is less stable than the wildtype. We also assessed the effect of the c.571_572delinsAG alteration on RNA-splicing using a minigene that contained exon 7 and 8 and their flanking introns. We found the c.571_572delinsAG substitution resulted in aberrant splicing of exon 8 (Figure 2B). The loss of exon 8 in the native *COCH* mRNA causes a shift in the reading frame resulting in a premature stop codon (p.Asp161Valfs4*). This mutant mRNA is expected to undergo NMD and be a null allele^29,30^. Interestingly, the spliced products of the constructs carrying the c.571_572delinsAG variant also produced a correctly spliced transcript that is identical to the control minigene (Figure 2B), in all three cell lines tested. These results could be due to the specific cell types used that would differ from the mechanism happening *in vivo* where the c.571_572delinsAG alteration would cause 100% exon skipping. Alternatively, c.571_572delinsAG could exert its effect via two mechanisms: aberrant splicing of exon 8, and protein misfolding.

In Proband B, we identified an ultra-rare, conserved, and predicted deleterious homozygous transversion variant c.271C>G (p.Arg91Gly). Although *in silico* prediction via HSF suggested this variant might alter RNA-splicing, we did not detect any differences in splicing between the wildtype and mutant minigenes (Supp Figure 1). The p.Arg91 residue is located in the LCCL domain (Figure 1E) and the significant change in folding energy strongly suggests this variant destabilizes the protein^23^. Besides the difference in charge between arginine and glycine, glycines are notoriously very flexible and can disturb the required rigidity of protein domains. This increase in flexibility fits the significant change in folding energy. Two variants in *LOXHD1* were also identified in Proband B. Pathogenic variants in LOXHD1 are responsible for deafness at the DFNB77 locus^32^. DFNB77-related deafness shows variability in onset, severity and progression^33–35^, including prelingual downsloping mild hearing loss. Based on phenotype alone we cannot rule out the involvement of *LOXHD1* in Proband B’s deafness. Given the reported consanguinity and conflicting *in silico* predictions for the *LOXHD1* variants, the *COCH* variant represents the most likely cause of hearing loss in this proband, however its disease-causing status remains uncertain and as such we classified it as variant of uncertain significance.

Finally, in Proband C, we detected a homozygous 9 nucleotide inframe deletion in exon 11. The deletion is predicted to remove the first amino acid from the vWFA2 domain and the two proceeding amino acids. We tested the effects of the 9-nucleotide deletion on splicing using a mini-genes containing wildtype and mutant exon 11. Spliced-products showed the loss of these nine nucleotides results in a transcript lacking exon 11 (Figure 2A). In the native mRNA, the loss of exon 11 shifts the reading frame leading to a truncated protein product (p.Ala321Glufs17*). We do not expect this allele to undergo the NMD pathway, given the newly created termination codon occurs in the last exon upstream of the native TAA. We do predict the translated protein to be nonfunctional, given the vWFA2 domain is missing along with ∼1/3 of the protein.

The auditory phenotype described in patients in this study is similar to the previously described hearing loss in DFNB110 patients, congenital downsloping mild/moderate-to-severe hearing loss. One subject homozygous for the p.Arg98Ter variant had vestibular dysfunction^2^. While not directly tested for, none of the patients in this study report balance problems or vertigo, similar to reported DFNB110 cases. Follow up studies are needed to see if patients with bi-allelic inactivating alleles of *COCH* also develop vestibular defects. Interestingly, the human phenotype is in sharp contrast to the auditory phenotype seen in the *Coch* knock-out mouse^26^. In the mouse, loss of *Coch* results in very mild and slightly progressive hearing impairment that is detectable after one year of age in the high frequencies^26^. These mice also exhibit elevated vestibular evoked potentials (VsEP) after 1 year of age, suggesting a potential vestibular defect, without any apparent menifestation of circling or twirling behavior, the hallmarks of vestibular impairment.

Our data highlight the need to exercise caution when classifying genetic variants in *COCH* as different pathologic mechanisms are involved, gain of function/dominant-negative for ADNSHL and loss- of-function for ARNSHL^36–40^. First, variants that result in a null allele should be considered likely pathogenic for DFNB110 and not DFNA9. Second, rare missense variants in *COCH* identified in patients with DFNB110 phenotype should be thoroughly investigated for their effect on RNA-splicing. If functional studies validate their deleterious effect on splicing, these variants should be considered likely pathogenic. The primary emphasis on coding variant interpretation focuses on the impact at the protein level. Often, if the variant falls outside the traditional splicing window, its impact on splicing is rarely considered. However, several studies have illustrated coding variant’s effect on splicing is underappreciated^13,14,41,42^.

In summary, we have expanded the mutational spectrum of *COCH*-related hearing loss to include missense, dinucleotide substitution, and inframe deletion variants. Importantly, our data further emphasize the importance of comprehensive variant interpretation irrespective of the conceived predicted translational impact as coding variants could also have a damaging impact on RNA-splicing.

## Acknowledgments

We would like to thank the patients for their participation in this study. This study was funded by NIDCDs R01’s DC002842, DC012049 and DC017955 to RJS and R56DC011803 to SR.

## Contributions

KTB, AG, MH, ZMA, RJS, HA and SR: conception and study design; KTB, MR, MH, MS, LTH, KF, EMR, NC performed experiments; KTB, AG, ZMA, RJH, HA, SR analyzed data and drafted the initial manuscript. All authors have reviewed and approved the finalized manuscript.

## Figures

**Supp Figure 1**. Minigene splicing assays. Gel electrophoresis and sequence chromatograms of wildtype, empty pET01 vector, and the c.271C>G mutating. Visualization and Sanger sequencing showed no difference in splicing between the wildtype and mutant.

**Supp Figure 2**. COCH ortholog aligment. Alignment of 11 COCH orthologs shows the conservation at residues: p.Arg91, p.Lys147, p.Val191, p.Glu211 and p.Ser365-Asn367 across species.

**Supp Table 1**. *LOXHD1* variants seen in Proband B.

## References

1. Azaiez, H. et al. Genomic Landscape and Mutational Signatures of Deafness-Associated Genes. Am. J. Hum. Genet. 103, 484–497 (2018).

2. Janssensdevarebeke, S. P. F. et al. Bi-allelic inactivating variants in the COCH gene cause autosomal recessive prelingual hearing impairment. Eur. J. Hum. Genet. 26, 587–591 (2018).

3. Mehregan, H. et al. Novel Mutations in KCNQ4, LHFPL5 and COCH Genes in Iranian Families with Hearing Impairment. Arch. Iran. Med. 22, 189–197 (2019).

4. Bae, S. H. et al. Identification of pathogenic mechanisms of COCH mutations, abolished cochlin secretion, and intracellular aggregate formation: Genotype-phenotype correlations in DFNA9 deafness and vestibular disorder. Hum. Mutat. 35, 1506–1513 (2014).

5. Robertson, N. G. et al. Isolation of novel and known genes from a human fetal cochlear cDNA library using subtractive hybridization and differential screening. Genomics 23, 42–50 (1994).

6. Robertson, N. G. et al. Mapping and characterization of a novel cochlear gene in human and in mouse: A positional candidate gene for a deafness disorder, DFNA9. Genomics 46, 345–354 (1997).

7. Bischoff, A. M. L. C. et al. Vestibular deterioration precedes hearing deterioration in the P51S COCH mutation (DFNA9): An analysis in 74 mutation carriers. Otol. Neurotol. 26, 918–925 (2005).

8. Kemperman, M. H. et al. DFNA9/COCH and its phenotype. Adv. Otorhinolaryngol. 61, 66–72 (2002).

9. Tsukada, K. et al. Detailed Hearing and Vestibular Profiles in the Patients with COCH Mutations. Ann. Otol. Rhinol. Laryngol. 124, 100S–110S (2015).

10. Hildebrand, M. S. et al. A novel mutation in COCH-implications for genotype-phenotype correlations in DFNA9 hearing loss. Laryngoscope 120, 2489–2493 (2010).

11. Booth, K. T. et al. PDZD7 and hearing loss: More than just a modifier. Am. J. Med. Genet. Part A 167, 2957–2965 (2015).

12. Booth, K. T. et al. Variants in CIB2 cause DFNB48 and not USH1J. Clin. Genet. 93, 812–821 (2018).

13. Booth, K. T., Kahrizi, K., Najmabadi, H., Azaiez, H. & Smith, R. J. Old gene, new phenotype: Splice-altering variants in CEACAM16 cause recessive non-syndromic hearing impairment. J. Med. Genet. 55, 555–560 (2018).

14. Booth, K. T. et al. Exonic mutations and exon skipping: Lessons learned from DFNA5. Hum. Mutat. 39, 433–440 (2018).

15. Richard, E. M. et al. Global genetic insight contributed by consanguineous Pakistani families segregating hearing loss. Hum. Mutat. 40, 53–72 (2019).

16. Zein, W. M. et al. Cone responses in usher syndrome types 1 and 2 by microvolt electroretinography. Investig. Ophthalmol. Vis. Sci. 56, 107–114 (2015).

17. Havrilla, J. M., Pedersen, B. S., Layer, R. M. & Quinlan, A. R. A map of constrained coding regions in the human genome. Doi.Org 51, 220814 (2017).

18. Liu, X., Wu, C., Li, C. & Boerwinkle, E. dbNSFP v3.0: A One-Stop Database of Functional Predictions and Annotations for Human Nonsynonymous and Splice-Site SNVs. Hum. Mutat. 37, 235–241 (2016).

19. Rentzsch, P., Witten, D., Cooper, G. M., Shendure, J. & Kircher, M. CADD: predicting the deleteriousness of variants throughout the human genome. Nucleic Acids Res. 47, D886–D894 (2019).

20. Desmet, F. O. et al. Human Splicing Finder: An online bioinformatics tool to predict splicing signals. Nucleic Acids Res. 37, 1–14 (2009).

21. Booth, K. T. et al. Splice-altering variant in COL11A1 as a cause of nonsyndromic hearing loss DFNA37. Genetics in Medicine 0, 1–7 (2018).

22. Venselaar, H., Te Beek, T. A. H., Kuipers, R. K. P., Hekkelman, M. L. & Vriend, G. Protein structure analysis of mutations causing inheritable diseases. An e-Science approach with life scientist friendly interfaces. BMC Bioinformatics 11, 548 (2010).

23. Quan, L., Lv, Q. & Zhang, Y. STRUM: Structure-based prediction of protein stability changes upon single-point mutation. Bioinformatics 32, 2936–2946 (2016).

24. Robertson, N. G. et al. Mutations in a novel cochlear gene cause DFNA9, a human nonsyndromic deafness with vestibular dysfunction. Nat. Genet. 20, 299–303 (1998).

25. Robertson, N. G. Inner ear localization of mRNA and protein products of COCH, mutated in the sensorineural deafness and vestibular disorder, DFNA9. Hum. Mol. Genet. 10, 2493–2500 (2001).

26. Jones, S. M. et al. Hearing and vestibular deficits in the Coch-/-null mouse model: Comparison to the CochG88E/G88E mouse and to DFNA9 hearing and balance disorder. Hear. Res. 272, 42–48 (2011).

27. Robertson, N. G. et al. Cochlin in normal middle ear and abnormal middle ear deposits in DFNA9 and Coch (G88E/G88E) mice. J. Assoc. Res. Otolaryngol. 15, 961–74 (2014).

28. Jung, J. et al. Cleaved Cochlin Sequesters Pseudomonas aeruginosa and Activates Innate Immunity in the Inner Ear. Cell Host Microbe 25, 513-525.e6 (2019).

29. Hentze, M. W. & Kulozik, A. E. A perfect message: RNA surveillance and nonsense-mediated decay. Cell 96, 307–310 (1999).

30. He, F. & Jacobson, A. Nonsense-Mediated mRNA Decay: Degradation of Defective Transcripts Is Only Part of the Story. Annu. Rev. Genet. 49, 339–366 (2015).

31. Makishima, T. et al. Targeted disruption of mouse Coch provides functional evidence that DFNA9 hearing loss is not a COCH haploinsufficiency disorder. Hum. Genet. 118, 29–34 (2005).

32. Grillet, N. et al. Mutations in LOXHD1, an Evolutionarily Conserved Stereociliary Protein, Disrupt Hair Cell Function in Mice and Cause Progressive Hearing Loss in Humans. Am. J. Hum. Genet. 85, 328–337 (2009).

33. Mori, K. et al. Mutations in LOXHD1 Gene Cause Various Types and Severities of Hearing Loss. Ann. Otol. Rhinol. Laryngol. 124, 135S–141S (2015).

34. Maekawa, K. et al. Mutational spectrum and clinical features of patients with LOXHD1 variants identified in an 8074 hearing loss patient cohort. Genes (Basel). 10, (2019).

35. Bai, X. et al. Five Novel Mutations in LOXHD1 Gene Were Identified to Cause Autosomal Recessive Nonsyndromic Hearing Loss in Four Chinese Families. Biomed Res. Int. 2020, (2020).

36. Azaiez, H. et al. TBC1D24 Mutation Causes Autosomal-Dominant Nonsyndromic Hearing Loss. Hum. Mutat. 35, 819–823 (2014).

37. Rehman, A. U. et al. Mutations in TBC1D24, a gene associated with epilepsy, also cause nonsyndromic deafness DFNB86. Am. J. Hum. Genet. 94, 144–152 (2014).

38. Mustapha, M. et al. An alpha-tectorin gene defect causes a newly identified autosomal recessive form of sensorineural pre-lingual non-syndromic deafness, DFNB21. Hum. Mol. Genet. 8, 409–12 (1999).

39. Verhoeven, K. et al. Mutations in the human alpha-tectorin gene cause autosomal dominant non-syndromic hearing impairment. Nat. Genet. 19, 60–2 (1998).

40. Vona, B., Nanda, I., Hofrichter, M. A. H., Shehata-Dieler, W. & Haaf, T. Non-syndromic hearing loss gene identification: A brief history and glimpse into the future. Mol. Cell. Probes 29, 260–270 (2015).

41. Collin, R. W. J. et al. Mid-frequency DFNA8/12 hearing loss caused by a synonymous TECTA mutation that affects an exonic splice enhancer. Eur. J. Hum. Genet. 16, 1430–6 (2008).

42. Aparisi, M. J. et al. Study of USH1 splicing variants through minigenes and transcript analysis from nasal epithelial cells. PLoS One 8, e57506 (2013).

