## Supplemental Table 1 for "Novel Loss-of-Function Mutations in *COCH* Cause Autosomal Recessive Nonsyndromic Deafness"

| **Variant** | **HGVSc** | **HGVSp** | **dbSNP** | **MAX gnomAD (%)** | **GERP** | **phyloP** | **SIFT** | **PolyPhen2** | **MutationTaster** | **LRT** | **CADD** |
| --- | --- | --- | --- | --- | --- | --- | --- | --- | --- | --- | --- |
| 18:g.44198212C>A | c.553G>T | p.Gly185Cys | rs758513799 | 0.003304 | 5.57 | 0.89 | D | B | D | - | 24 |
| 18:g.44104434C>G | c.4871G>C | p.Gly1624Ala | rs757775256 | 0.03514 | 5.06 | 0.85 | T | P | D | N | 21.5 |

**Supplementary Table 1**

***LOXHD1* variants identified in Proband B**. Nucleotide numbering: the A of the ATG translation initiation site is noted as +1 using transcript NM_144612.6 of *LOXHD1*. D, predicted Damaging or Deleterious; −, data not available; T, Tolerated; A, disease automatic; B, Benign; P, Possibly Damaging; N, Neutral.
