## Supplementary figures and images for "Novel Loss-of-Function Mutations in *COCH* Cause Autosomal Recessive Nonsyndromic Deafness"

### Supplemental figure 1

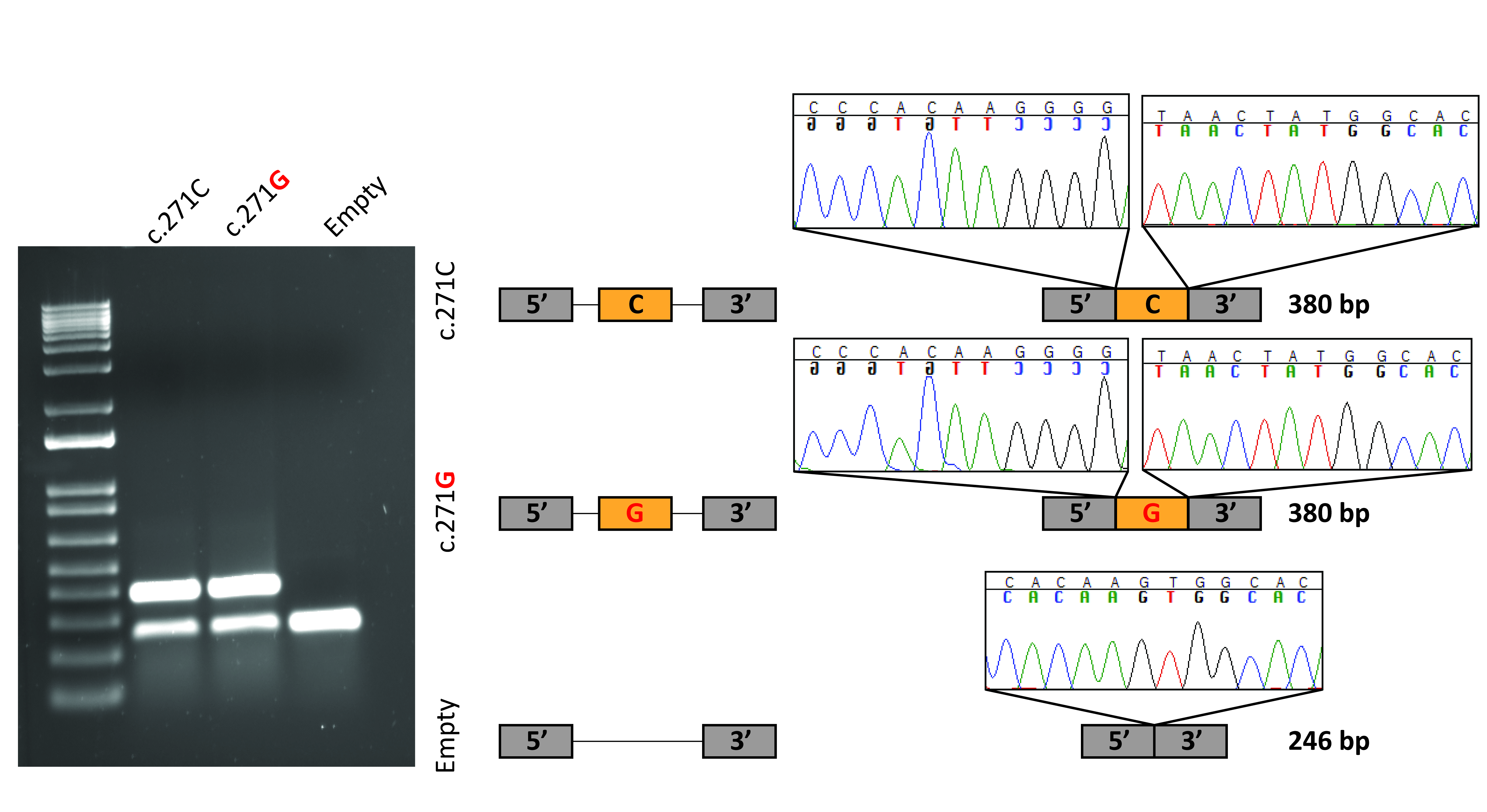

### Supplemental Figure 2

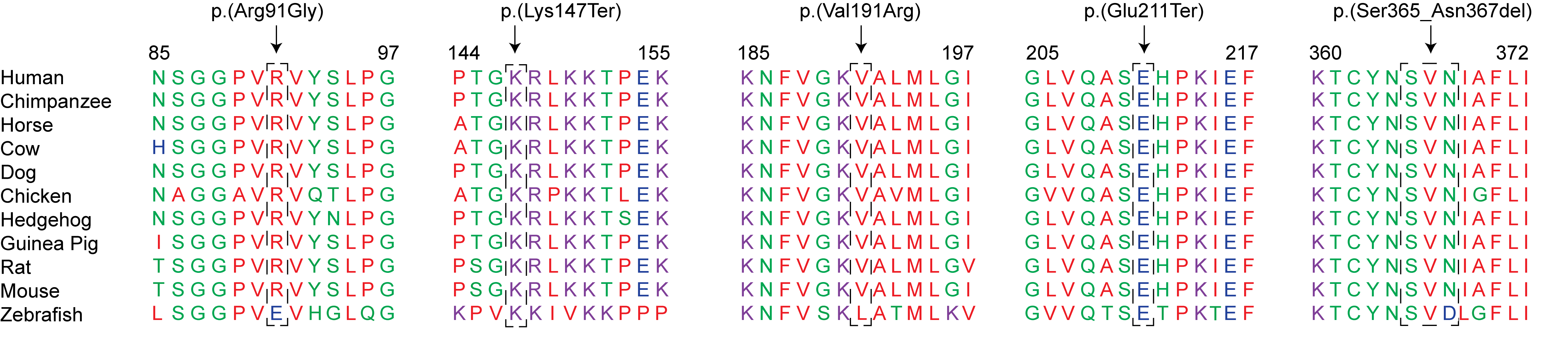
